## Supplementary Figures S1-S12 for "Removal of developmentally regulated microexons has a minimal impact on larval zebrafish brain morphology and function"

Microexon and surrounding exons A-N

Zoom of gRNAS/UGC/UC A-N

Microexon and surrounding exons P-V

Zoom of gRNAS/UGC/UC P-V

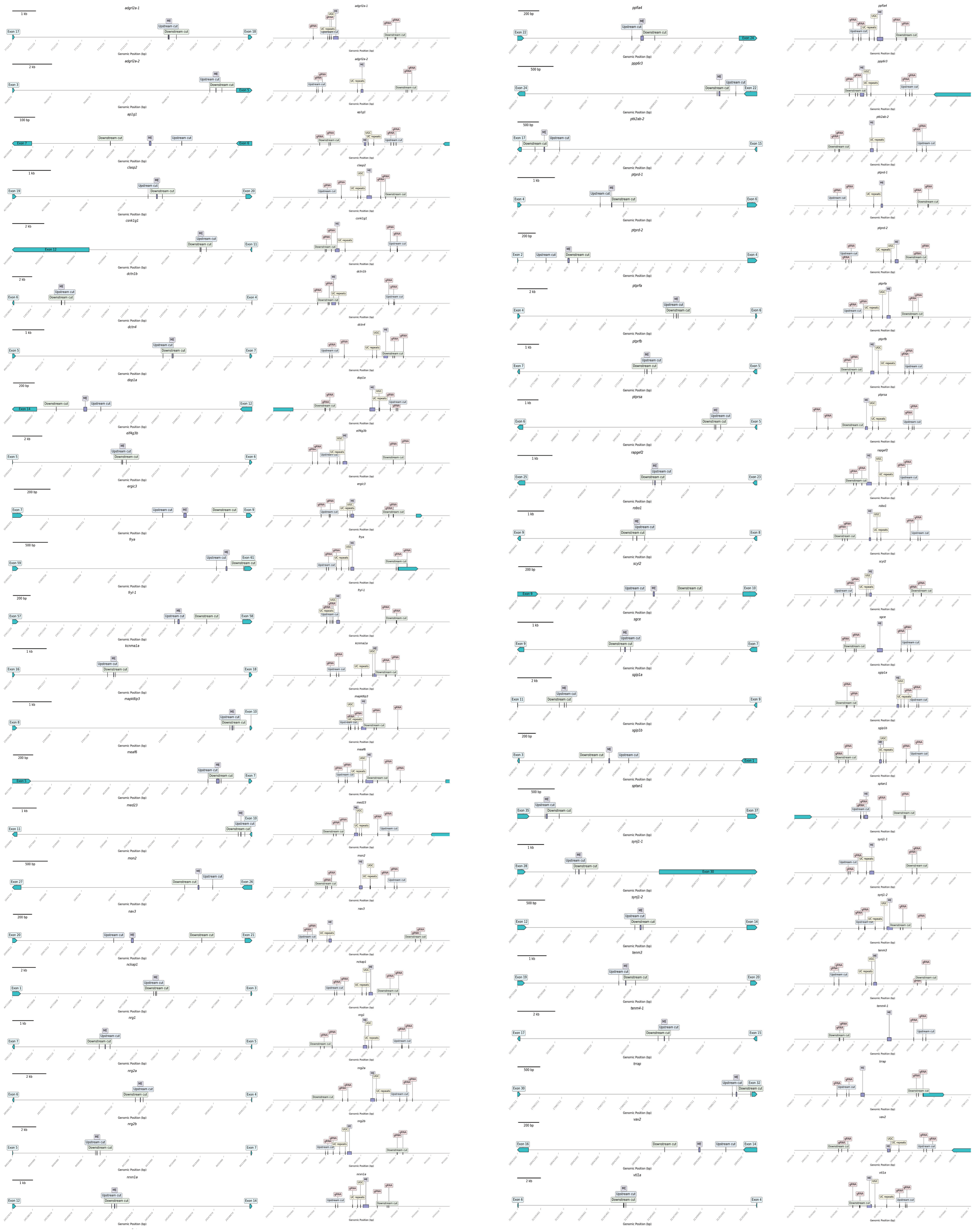

**Figure S1. Microexons in relationship to cut sites, neighboring exons, putative regulatory elements, and gRNA sites.** (**A**) Diagrams were generated with DNA Features Viewer (https://edinburgh-genome-foundry.github.io/DnaFeaturesViewer/). The first and third columns show the microexons and neighboring exons, with a unique scale bar included for each mutant. The second and fourth columns show the zoomed area around each microexon with the start position for putative regulatory elements and gRNA sites. When the microexon neighbored either the first or last exon, the UTR is included in the labeled exon (e.g., *csnk1g1* and *synj1-1*). When determining possible UC-repeat elements, a distance of 100 base-pairs from the microexon was considered although <50 base-pairs is canonical. The label for the UC repeat represents the beginning of possible repeat sequences and not the span; thus, there may be closer elements.

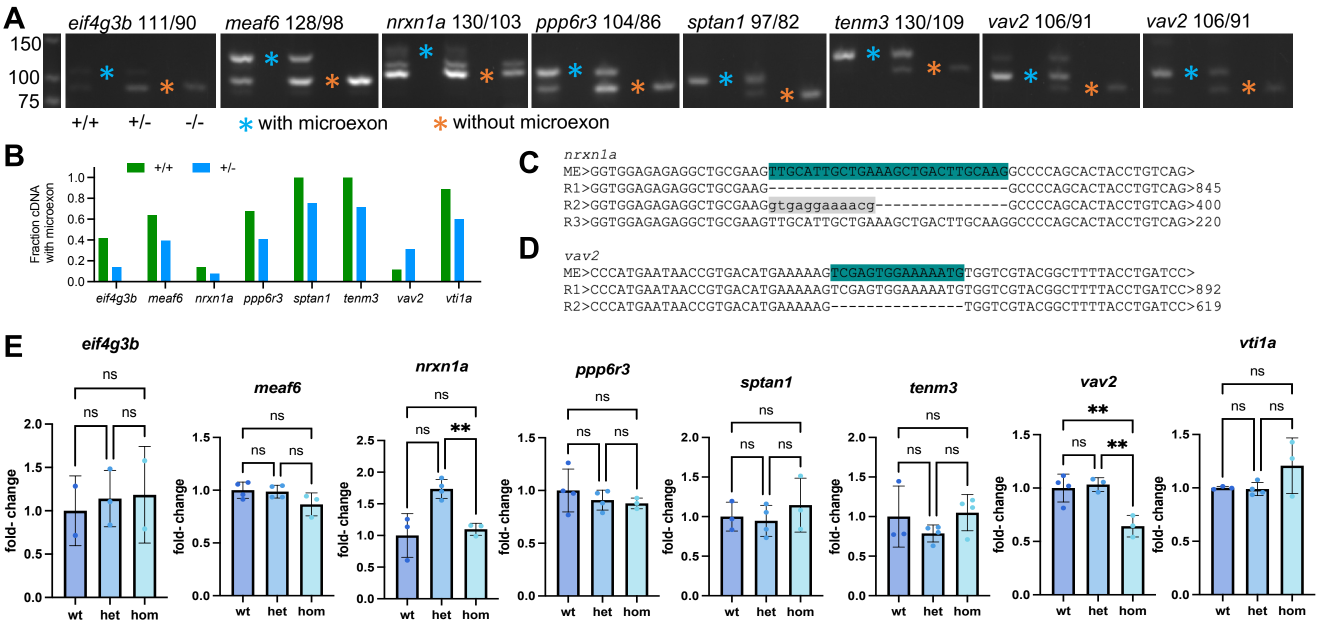


**Figure S2. Transcriptomic outcomes of eliminating selected microexons.** (**A**) RT-PCR validations of microexon removal at 6 dpf for wildtype (+/+), heterozygous (+/-) and homozygous (-/-) siblings. The expected length of the product with and without the microexon are shown above each sample, and the ladder is shown on the left side. The upper band in the *vav2* wild-type and heterozygous samples is a heterodimer of the two products, an outcome we often see when genotyping smaller deletions. (**B**) Quantification of the gels in panel A. (**C**) Nanopore sequencing (Plasmidsaurus) of the heterozygous *nrxn1a* sample with corresponding read counts (right). The second read (R2) represents the intermediate band on the *nrxn1a* gel above. This sequence corresponds to an extension of the upstream exon (gray), indicating that there are two isoforms of *nrxn1a* in addition to the developmentally regulated inclusion of the microexon. (**D**) Nanopore sequencing (Plasmidsaurus) of the heterozygous *vav2* sample with corresponding read counts. The R2 sequencing missing the microexon was additionally confirmed with Sanger sequencing of the homozygous sample. (**E**) qRT-PCR for selected microexon lines. The expression of each gene was normalized to *rpl13a*. Statistical significance was calculated with the Brown-Forsythe and Welch ANOVA corrected for multiple comparisons.

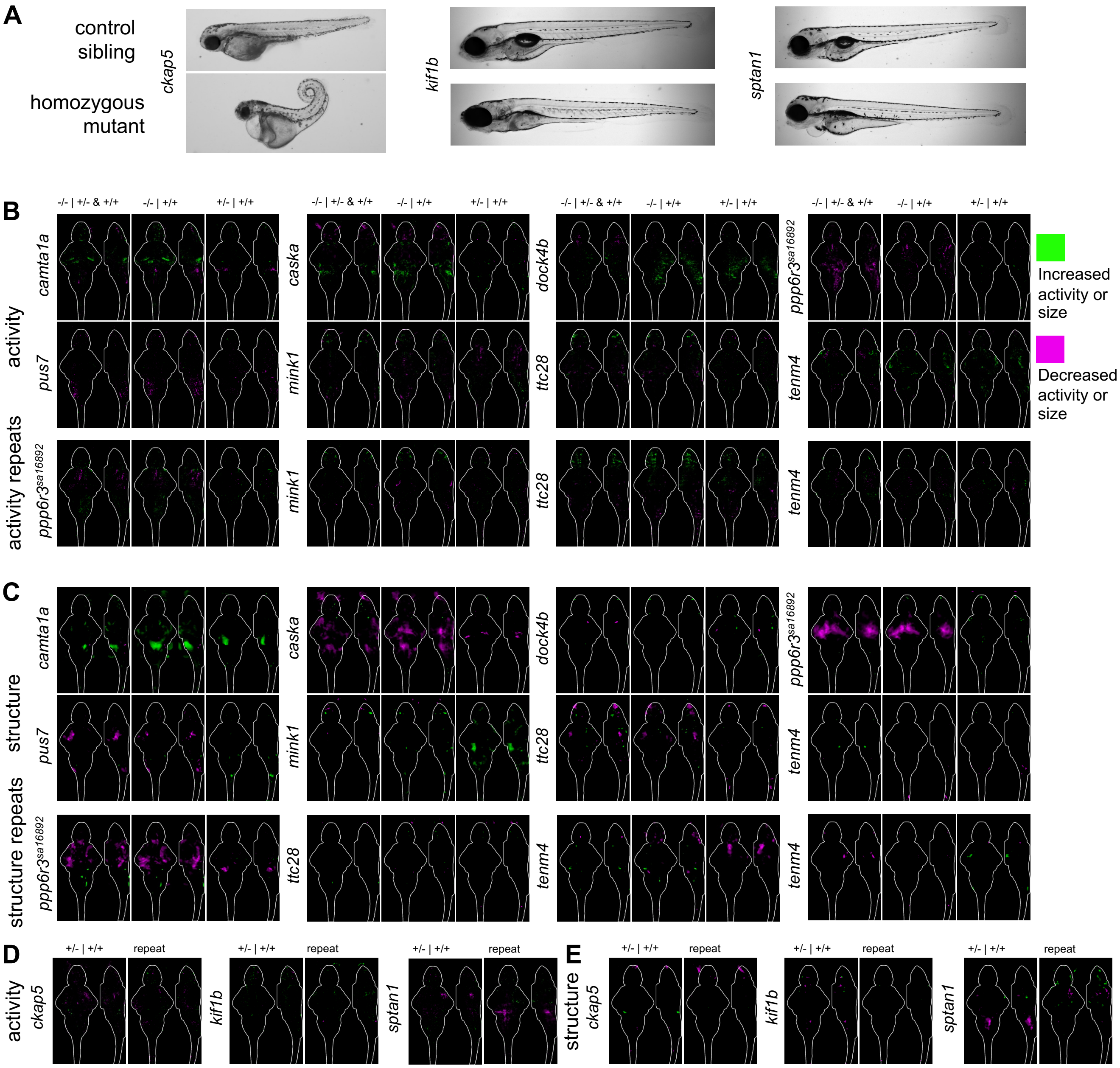


**Figure S3. Phenotypes for mutants in microexon genes that remove protein sequence beyond the microexon.** (**A**) Developmental phenotypes observed in homozygous larvae for three genes with truncating mutations. Chi Square analysis $P < 0.01$ from 12/12 animals with and without a phenotype. Additionally, healthy homozygous animals were never recovered from multiple heterozygous incrosses assessed at 4-6 dpf. (**B**) Brain activity maps for heterozygous incrosses. The comparisons are shown above the sum-of-slices intensity projection with a 6 dpf brain outline, where the genotype before the | is being compared to the one after. All N are in Table S2. (**C**) Brain structure maps for heterozygous incrosses. All N are in Table S2. (**D**) Brain activity maps for heterozygous outcrosses for those mutants with lethal developmental phenotypes that prohibit homozygous imaging. The comparisons are shown above the sum-of-slices intensity projection with a 6 dpf brain outline, where the genotype before the | is being compared to the one after. All N are in Table S2. (**E**) Brain structure maps for heterozygous outcrosses for those mutants with lethal developmental phenotypes that prohibit homozygous imaging. The comparisons are shown above the sum-of-slices intensity projection with a 6 dpf brain outline, where the genotype before the | is being compared to the one after. All N are in Table S2.

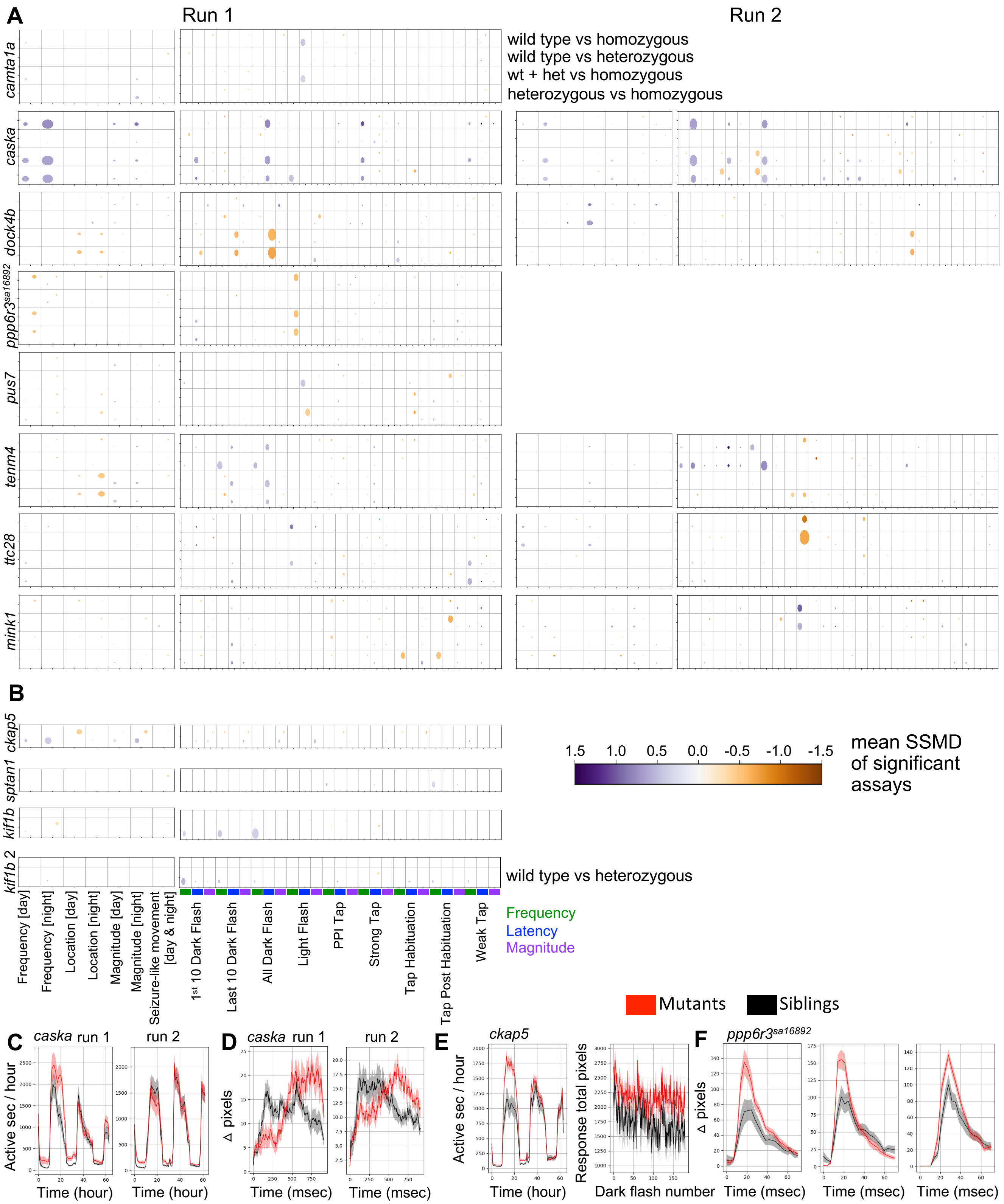


**Figure S4. Behavior summary data for mutants in microexon genes that remove protein sequence beyond the microexon.** (**A**) Baseline and stimulus-driven behavior dot plots for heterozygous incrosses. Dot size corresponds to the percent of significant assays in the category (e.g., Magnitude). Replicate experiments using different parental pairs are shown side-by-side. All N are in Table S2. (**B**) Baseline and stimulus-driven behavior dot plots for heterozygous outcrosses. Dot size corresponds to the percent of significant assays in the category (e.g., Magnitude). (**C**) The *caska* truncating mutant has repeatable phenotypes in motion frequency and dark flash response Example frequency of motion plot for *caska* mutants. The nighttime movement frequency is increased. The wild-type siblings (black) are compared to the homozygous (red). (**D**) Graph demonstrating the reduced dark flash responsivity of *caska* mutants. The latency is increased in this line (see panel H), which can be visualized by this response plot. (**E**) Increased daytime movement frequency and increased dark flash response frequency in *ckap5* heterozygous mutants (red) compared to wild-type siblings (black). This phenotype represents the strongest behavioral outcome for heterozygous compared to wild type. (**F**) The *ppp6r3*$^{sa16892}$ has an increased response to acoustic stimuli. These three plots are response traces (shown from left to right) induced by strong taps that occur after tap habituation block 2 (day5dpfhab2post), after tap habituation block 3 (day5dpfhab3post), and during the night (a0f1000d5pD300a1f1000d5p).

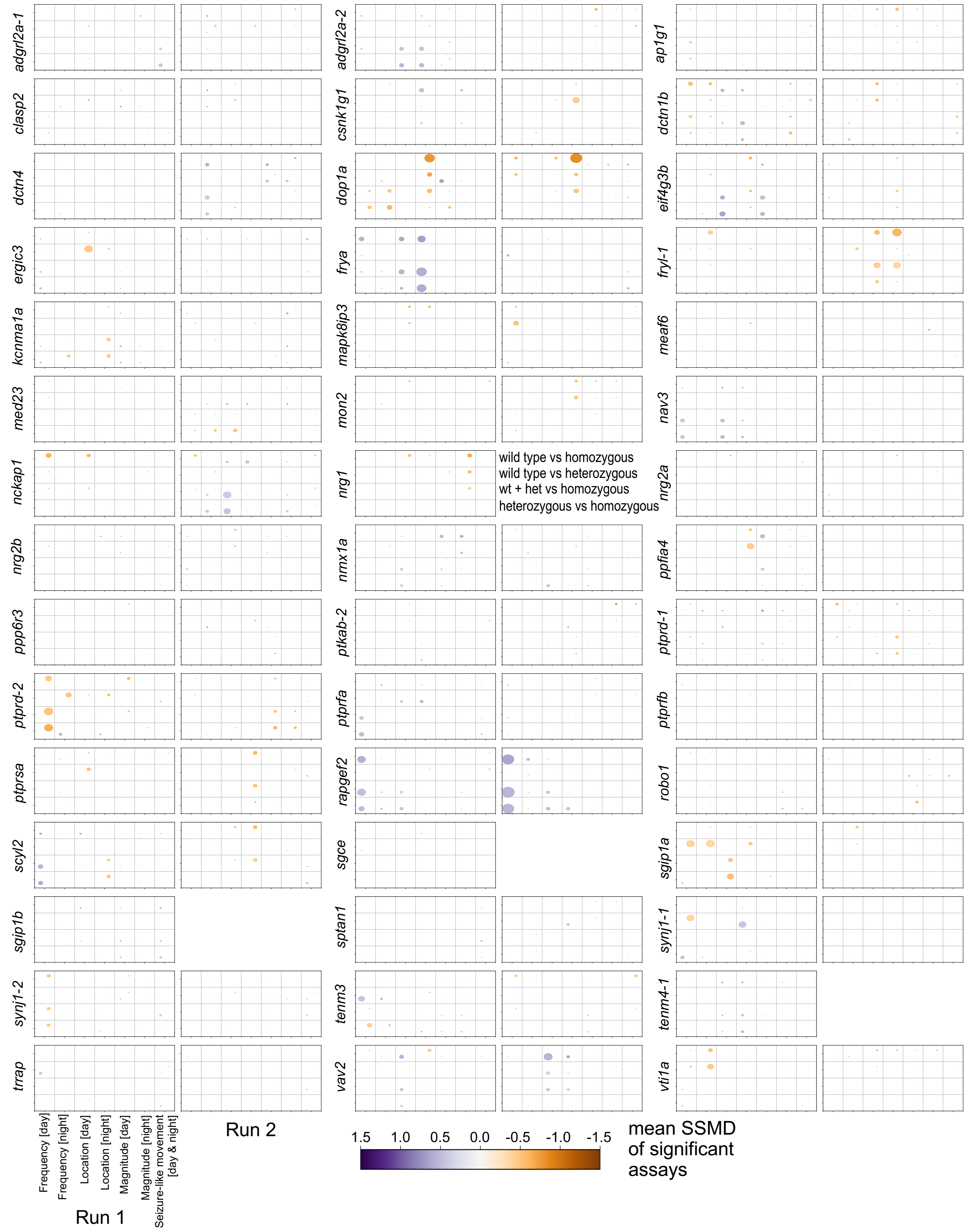


**Figure S5. Baseline behavior summary data for microexon mutants.** Baseline behavior dot plots for all sibling comparisons. Dot size corresponds to the percent of significant assays in the category (e.g., Magnitude). Replicate experiments using different parental pairs are shown side-by-side. All N are in Table S2.

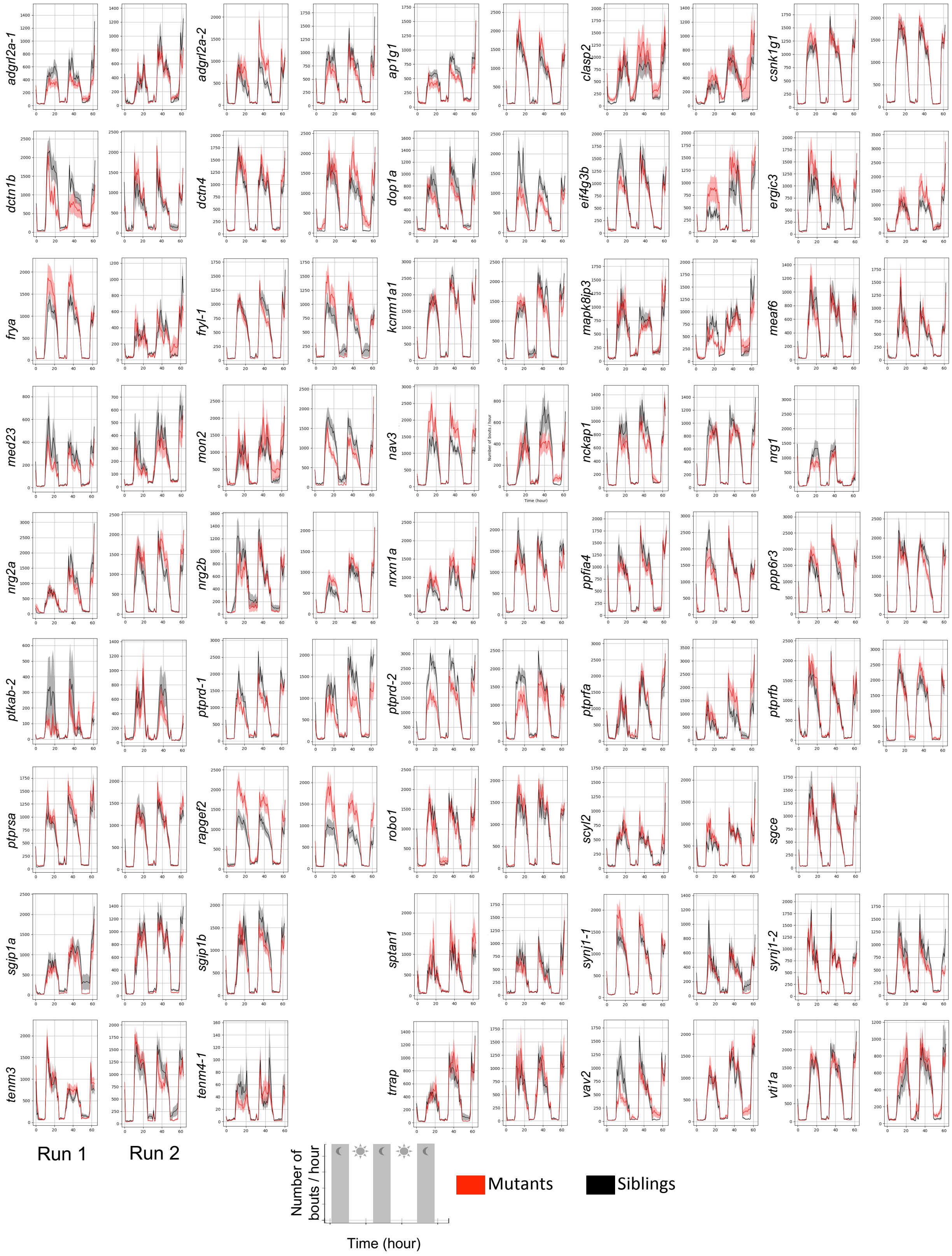


**Figure S6. Frequency of motion example plots for microexon mutants.** Frequency of motion measured as number of bouts/hour over the entire duration of the experiment. The biological replicates are shown side-by-side. All N are in Table S2. The wild-type siblings (black) are compared to the homozygous (red).

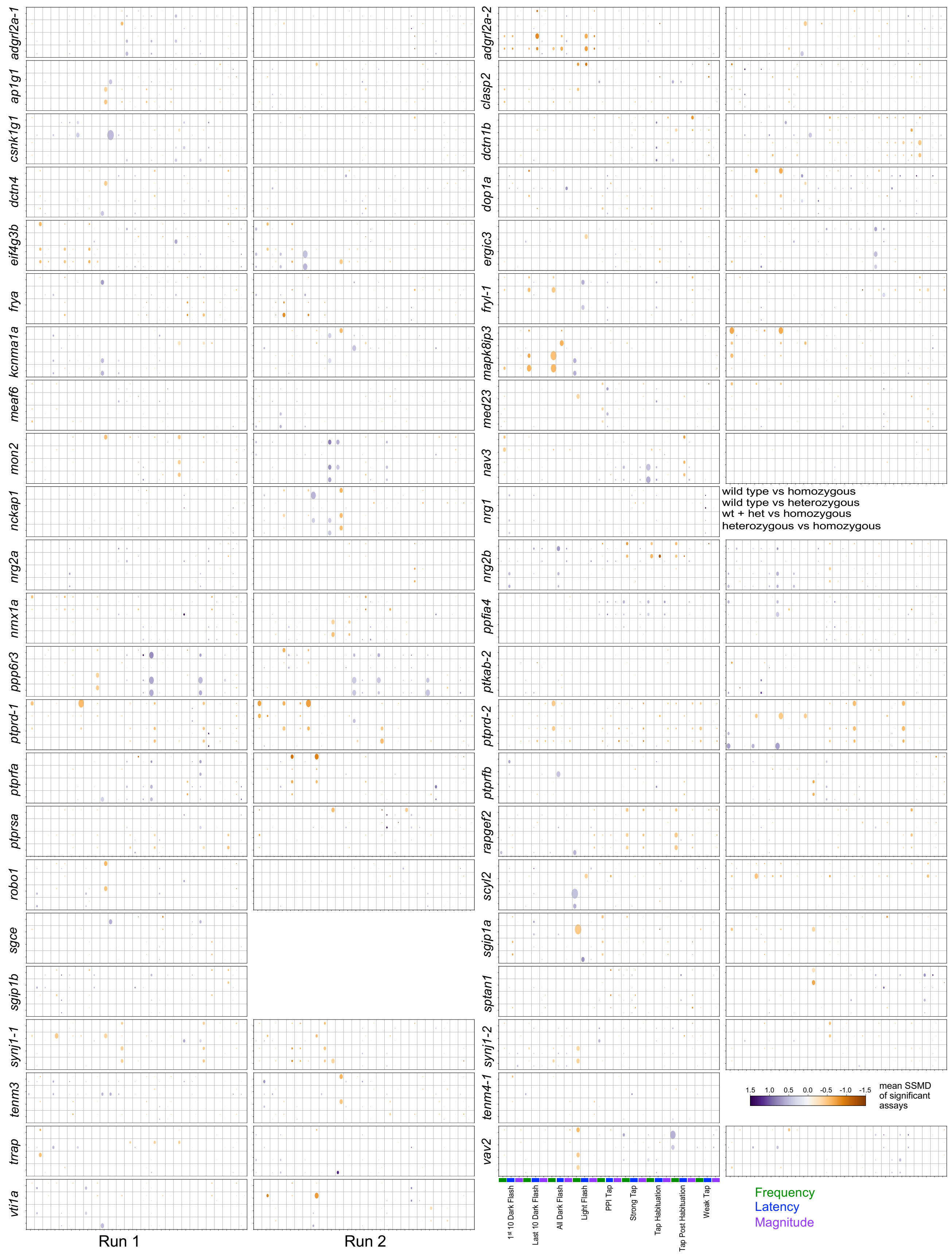


**Figure S7. Stimulus-driven behavior summary data for microexon mutants.** Stimulus-driven behavior dot plots for all sibling comparisons from a heterozygous parental in-cross.

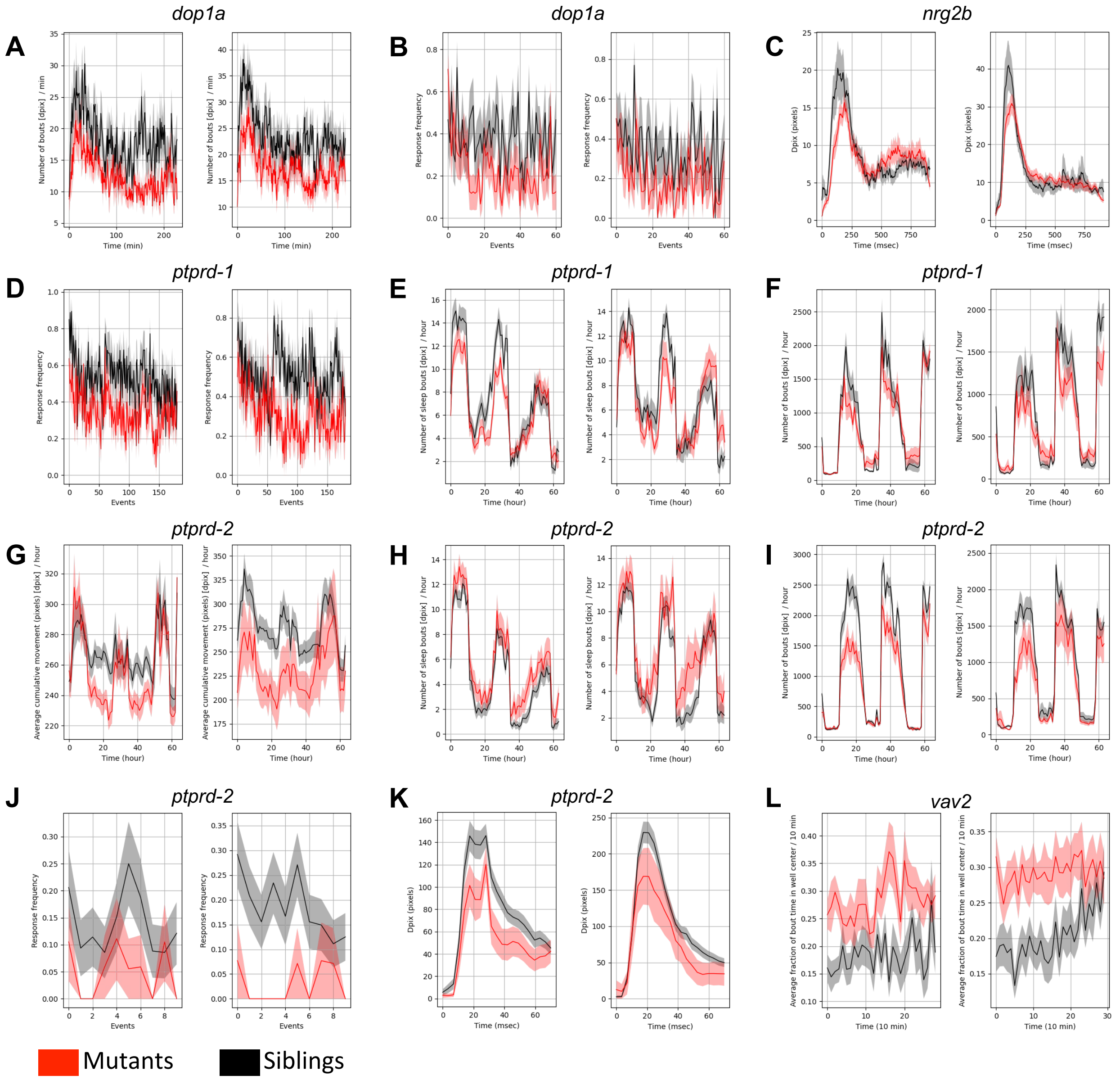


**Figure S8. Additional behavioral phenotypes for microexon mutants.** All N are in Table S2. (**A**) Frequency phenotype of *dop1a* mutants. Shown is the day3msdf_dpix_numberofbouts_60 graph for run 1/run 2. P-values: 0.033/0.038 (linear mixed model) and 0.036/0.11 (Kruskal-Wallis ANOVA, which performs poorly on data with a large range). (**B**) Reduced dark flash response frequency phenotype of *dop1a* mutants. Shown is the response frequency for dark flash block 3 (day6dpfdf3_responsefrequency). P-values: 0.047/0.036 (Kruskal-Wallis ANOVA). (**C**) Reduced dark flash response of *nrg2b* mutants. P-values are not calculated for response graphs. (**D**) Reduced dark flash response frequency phenotype of *ptprd*-1 mutants. Shown is the response frequency for dark flashes from all three blocks (day6dpfdfall_responsefrequency). P-values: 0.0056/0.0018 (Kruskal-Wallis ANOVA). (**E**) Reduced nighttime sleep bout frequency phenotype of *ptprd*-1 mutants. Shown is the plot for the entire 3-night/day experiment (combo plot). P-values: 0.001/0.013 (linear mixed model). The same measure from a subsection of the data, the entire second night (day2nightall_dpix_numberofboutsSLEEP, not shown), has Kruskal-Wallis ANOVA p-values of 0.001/0.003. (**F**) Altered number of bouts frequency phenotype of *ptprd*-1 mutants. Shown is the entire 3-night/day experiment (combo plot). P-values are not significant for the entire duration, but a subsection (day1morn_dpix_numberofbouts_60, not shown) has Kruskal-Wallis ANOVA p-values of 0.003/0.012. (**G**) Reduced daytime bout total pixels magnitude (dpix_boutcumulativemovement) phenotype of *ptprd*-2 mutants. Shown is the entire 3-night/day experiment (combo plot). P-values are not significant for this duration. The same measure from a subsection of the data, the evening of day 1 of the experiment (day1evening_dpix_boutcumulativemovement, not shown), has Kruskal-Wallis ANOVA p-values of 0.002/0.019. (**H**) Increased sleep bout (stronger in daytime) frequency phenotype of *ptprd*-2 mutants. Shown is the entire 3-night/day experiment (combo plot). P-values: 0.001/0.013 (Kruskal-Wallis ANOVA). The same measure from a subsection of the data, the entire second night (day2nightall_dpix_numberofboutsSLEEP_60, not shown), has Kruskal-Wallis ANOVA p-values of 0.001/0.01. (**I**) Reduced number of bouts (stronger in daytime) frequency phenotype of *ptprd*-2 mutants. Shown is the entire 3-night/day experiment (combo plot). P-values: 0.001/0.04 (linear mixed model) and 0.003/0.08 (Kruskal-Wallis ANOVA). (**J**) Reduced acoustic response frequency phenotype of *ptprd*-2 mutants. Shown are the escape responses for the strong stimuli occurring following acoustic habituation (habituation_day5dpfhab1post_responsefrequency). The responses are filtered for those that are true escapes, and the phenotype emerges for only the true, short-latency C-bends. P-value: 0.03/0.006 (Kruskal-Wallis ANOVA). (**K**) Reduced strong acoustic responses of *ptprd*-2 mutants. This response plot goes along with the frequency data, but this plot is not filtered for only the true C-bends. P-values are not calculated for response graphs (**L**) Increased daytime location center preference for *vav2* mutants. Shown is the evening of the second day (day2evening_boutcenterfraction), but multiple subsections show significant increase in center dwelling preference. P-value: 0.006/0.013 (Kruskal-Wallis ANOVA).

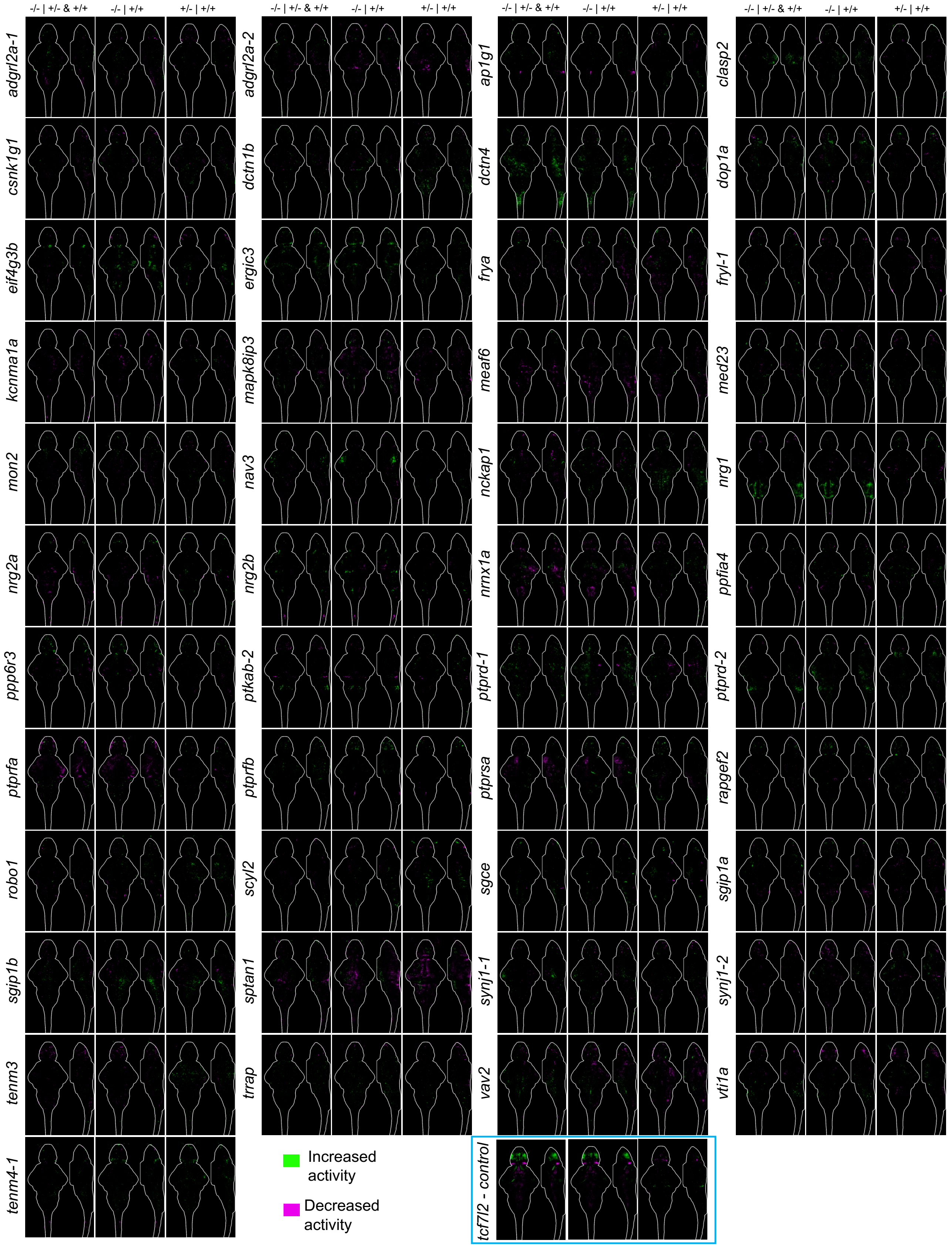


**Figure S9. Brain activity maps for multiple genetic comparisons for microexon mutants.** The comparisons are shown above the sum-of-slices intensity projection with a 6 dpf brain outline, where the genotype before the | is being compared to the one after. All N are in Table S2.

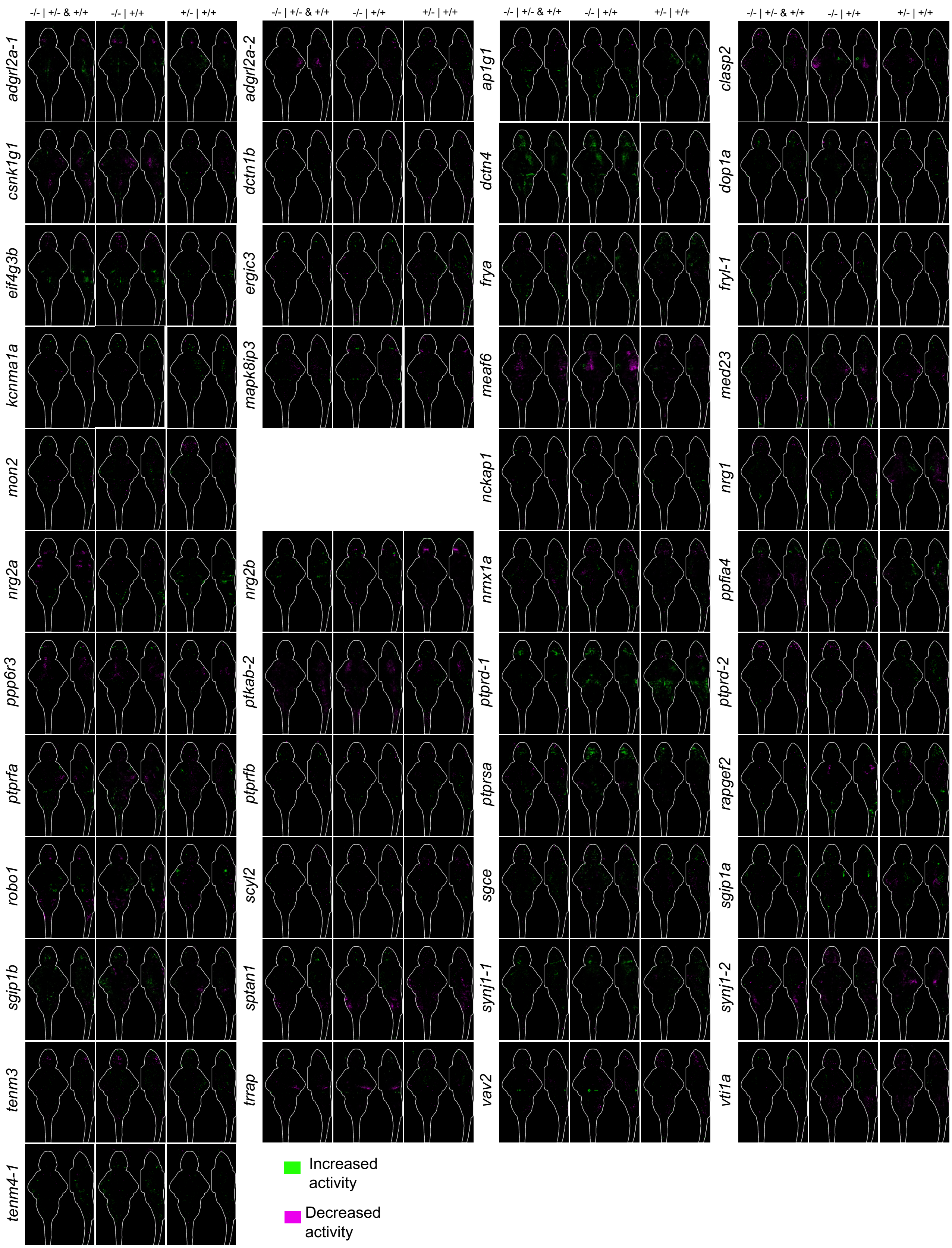


**Figure S10. Second set of brain activity maps for multiple genetic comparisons for microexon mutants.** The comparisons are shown above the sum-of-slices intensity projection with a 6 dpf brain outline, where the genotype before the | is being compared to the one after. All N are in Table S2.

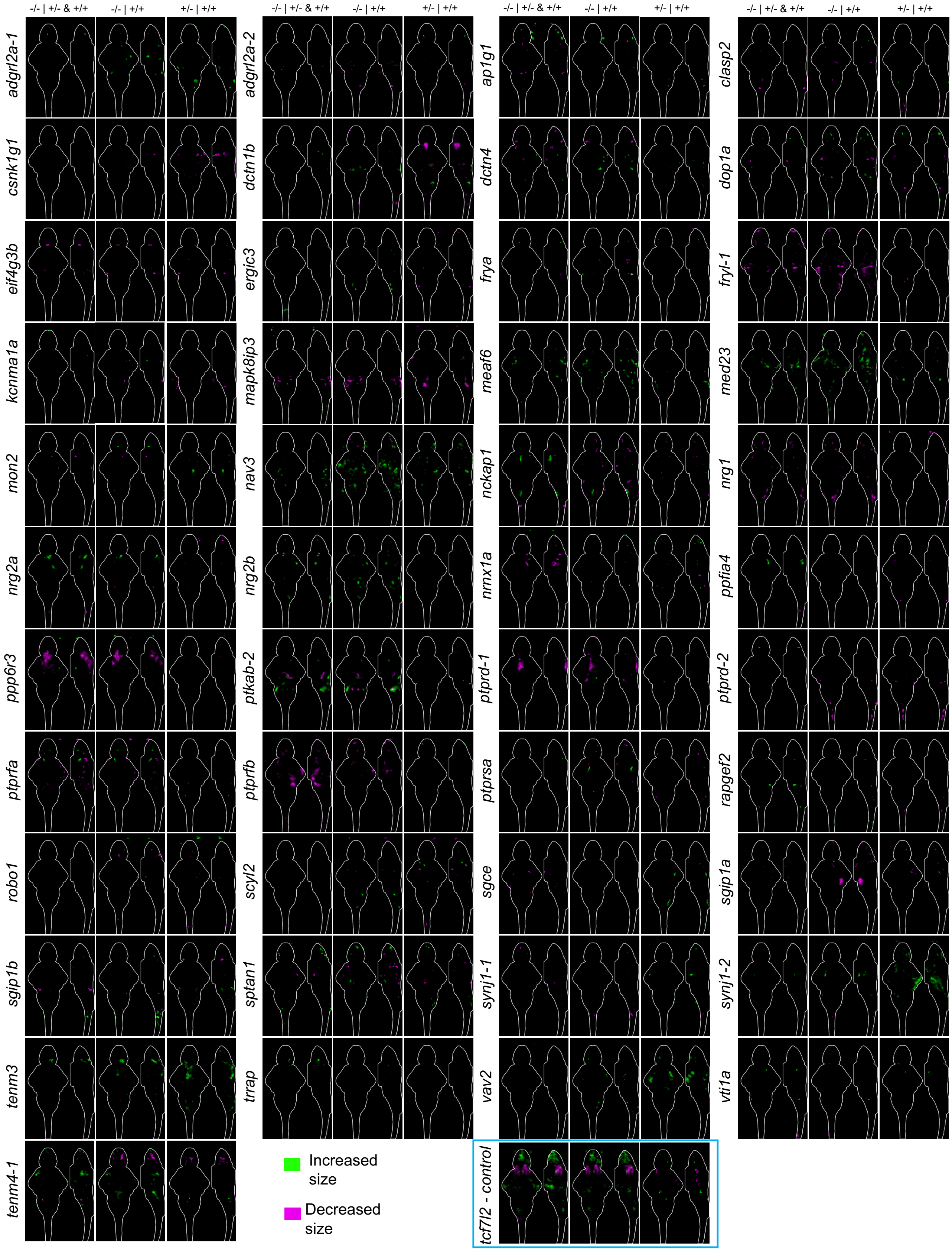


**Figure S11. Brain structural maps for multiple genetic comparisons for microexon mutants.** The comparisons are shown above the sum-of-slices intensity projection with a 6 dpf brain outline, where the genotype before the | is being compared to the one after. All N are in Table S2.

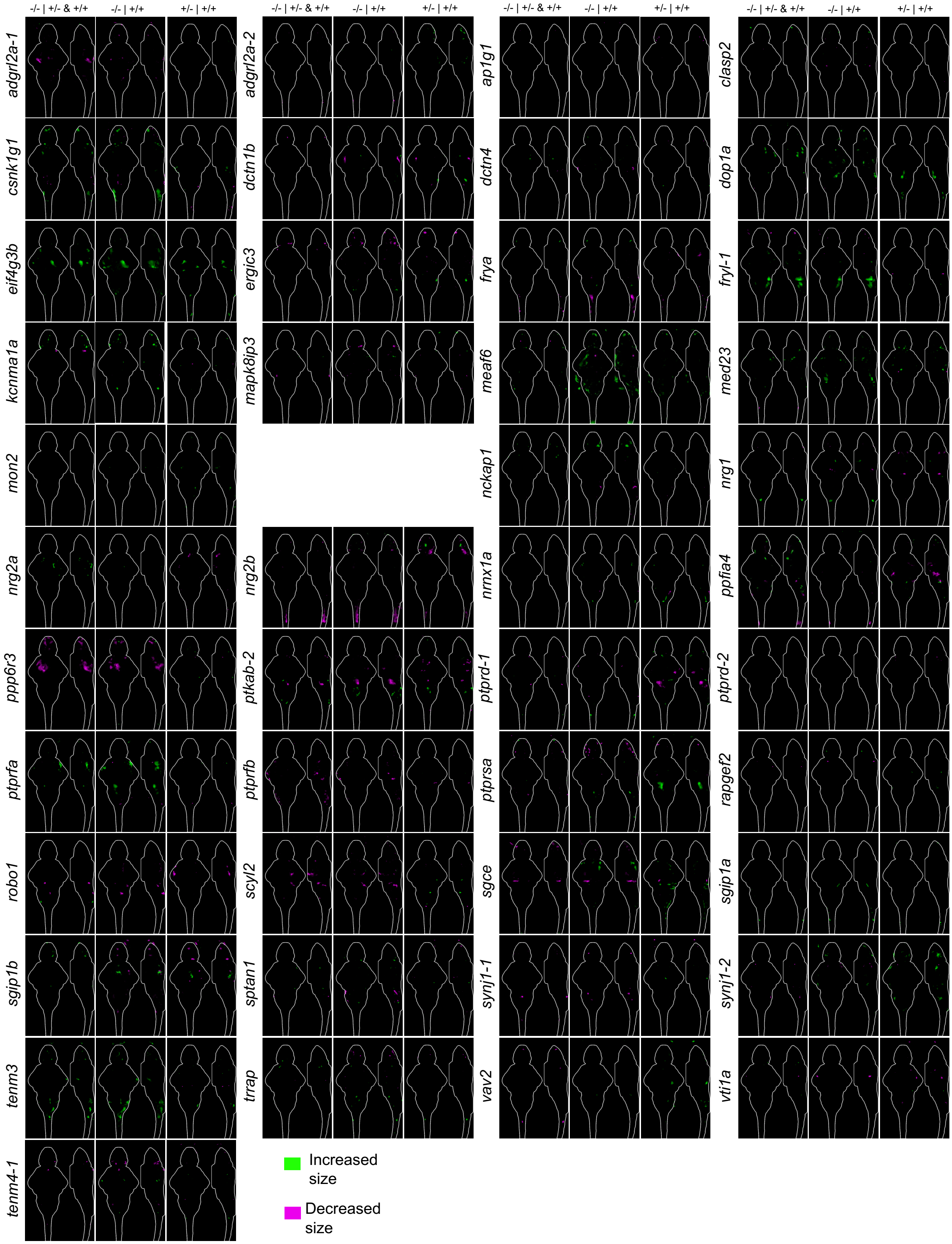


**Figure S12. Second set of brain structural maps for multiple genetic comparisons for microexon mutants.** The comparisons are shown above the sum-of-slices intensity projection with a 6 dpf brain outline, where the genotype before the | is being compared to the one after. All N are in Table S2.

**Table S1. Sequences and locations of 95 microexons conserved between zebrafish and mouse.**

**Table S2. Mutants generated and corresponding genotyping and experimental information.**
